# A tandem segmentation-classification approach for the localization of morphological predictors of *C. elegans* lifespan and motility

**DOI:** 10.1101/2021.05.16.444281

**Authors:** Artur Yakimovich, Evgeniy Galimov

**Affiliations:** Artificial Intelligence for Life Sciences CIC, London, United Kingdom

**Keywords:** CNN, Caenorhabditis elegans, image segmentation, image classification, aging, lifespan, deep learning, interpretable machine learning

## Abstract

*C. elegans* is an established model organism for studying genetic and drug effects on ageing, many of which are conserved in humans. It is also an important model for basic research, and *C. elegans* pathologies is a new emerging field. Here we develop a proof-of-principal convolutional neural network-based platform to segment *C. elegans* and extract features that might be useful for lifespan prediction. We use a dataset of 734 worms tracked throughout their lifespan and classify worms into long-lived and short-lived. We designed WormNet - a convolutional neural network (CNN) to predict the worm lifespan class based on young adult images (day 1 – day 3 old adults) and showed that WormNet, as well as, InceptionV3 CNN can successfully classify lifespan. Based on U-Net architecture we develop HydraNet CNNs which allow segmenting worms accurately into anterior, mid-body and posterior parts. We combine HydraNet segmentation, WormNet prediction and the class activation map approach to determine the segments most important for lifespan classification. Such a tandem segmentation-classification approach shows posterior part of the worm might be more important for classifying long-lived worms. Our approach can be useful for the acceleration of anti-ageing drug discovery and for studying *C. elegans* pathologies.

## Introduction

The nematode *Caenorhabditis elegans* (*C. elegans*) is an established model for studying various interventions into the ageing process, which allowed to find numerous genes and drugs interfering with aging. 5 out of 7 Tier 1 and 4 out of 6 Tier 2 anti-ageing drugs considered for human trials extend lifespan in the *C. elegans* model. There are many ageing pathways conserved among species and the worms are expected to be used extensively not only in longevity research but also in the appearing anti-ageing industry (1). Additionally, humanized worms are now used to establish promising models for neurodegeneration (2). However, unlike genetics of longevity, *C. elegans* phenotypes of ageing are not well studied yet. Particularly, we know little about age-related pathologies and their development, as well as, which pathologies determine lifespan and how they cause death (3). Several pathologies including gut atrophy, uterine tumours and pharyngeal infection were described recently (4–6). In this light, discovering new *C. elegans* pathologies, particularly determining lifespan, is becoming an important challenge. Studying pathologies in *C. elegans* might help to get a better understanding of the ageing process, as well as, the mechanisms and effects of anti-ageing drugs.

Recent advances in machine learning (ML) and deep learning (DL) (7) may aid aging studies employing *C. elegans* through uncovering and summarising previously unseen behavioural and morphological patterns in large experimental datasets. For example, in a recent work several physiological parameters were measured longitudinally and application of support vector regression allowed to explain different amount of variance in *C. elegans* lifespan by: movement (57%), cross-sectional size (5%), texture (42%), autofluorescence (52%), oocyte laying rate (28%) (8). Interestingly, it was found that the brood size correlates with lifespan in mated hermaphrodite (r = 0.28) (9). Furthermore, independent studies confirm that the muscle function is probably the best predicting physiological feature: fast pharyngeal pumping span (r = 0.49), and pharyngeal pumping span (r = 0.83) were found to be highly correlated with the lifespan length (Huang et al., 2004). Also, maximum velocity at day 9 (10) and the rate of speed decay (days 3-9) (11) explain 71% and 91% of variability in lifespan accordingly. Cellular and molecular predictors of *C. elegans* lifespan length were also discovered. Expression of hsp-16.2 induced by heat shock in day 1 adults was found to be correlated with lifespan (12). Free of confounding effects of interventions like heat shock, basal expression of sod-3 at day 9 also correlated with lifespan (r = 0.57), which probably reflects response to pathogenic food (13). mir-71 expression from day 4 onwards can be highly predictive and explains 47% of variability in lifespan (14). Strikingly, a strong inverse correlation (r = −0.93) between nucleolar size (measured on day 1) and longevity indicates deregulated protein synthesis as an important component of aging (15). Noteworthy, early on a Machine Vision approach was also applied to classify ageing phenotypes in *C. elegans*. Particularly, linear discriminant classifier was used to segregate images of pharynxes of different ages for subsequent molecular characterization (16).

Among other methods, one of the most powerful machine learning approaches, particularly for image analyses, is the use of convolutional neural networks (CNN) (7), which are inspired by visual cortex neural network organization. CNN allowed to achieve impressive results in image recognition, with near human performance on MNIST dataset and outperformed humans on a traffic sign recognition by a factor of two (17). CNN repeatedly showed best performance during “The ImageNet Large Scale Visual Recognition Challenge” in image classification (18, 19). Introduction of skipped connections to CNN dramatically improved their speed and accuracy, and such residual CNNs are now state-of-the-art for image classification (20, 21). Encoder-decoder residual networks like U-Net (22), V-Net and Tiramisu also outperform the classical boundary extraction, threshold and region-based methods used in medical image segmentation field (23). Despite the impressive results with DL approaches, one of the main drawbacks is that DL networks are black boxes so it is difficult to get the features important for decision making by the network (24). To circumvent this shortcoming, several saliency techniques have been proposed (25–27). One such technique is using the global average pooling layer to produce a so-called class activation map (CAM) and localize class-specific image regions in unsupervised manner (28). The produced generic localizable deep features can aid researchers in understanding the basis of discrimination used by CNNs for their tasks. However, thus far, no approaches to combine biologically meaningful image segmentation and classification saliency to facilitate phenotype discovery through interpretation have been developed.

Remarkably, CNN were recently used to predict lifespan in worms. In the first paper, a dataset of 913 images of *C. elegans* were used. Each timepoint (day) has at least 30 worms, and all of them were anesthetized before imaging. InceptionResNetV2-based architecture achieved a mean absolute error (MAE) of 0.96 day in the regression mode, and the accuracy of 57.6% in classification mode (29). In another work, the authors used automatic imaging system capable of tracking the same worm during the whole lifespan, so they had data for 734 worms for which images were taken every 3.5 hours. They used U-Net to segment worms from background and then preformed the worm body coordinate regression to create straightened worm representations. Then they used a modified ResNet34 and managed to regress worm age with minimal MAE of 0.6 days for raw images (30).

Here we used the same dataset as in (8, 30), however instead of predicting age of each worm, we develop a CNN-based platform we called WormNet capable of classifying young adults (day 1-3) into short-lived and long-lived, and also design an approach for extracting features important for such classification. Similarly, we have applied WormNet to classify *C. elegans* movement. To interpret classification results in a by-design fashion, we have companied classification CNN with a tandem segmentation CNN. For this, we devised a new U-Net-based architecture (HydraNet) for segmenting worms from the background and also segmenting worm’s body into anterior, mid-body and posterior parts. Interpretation of the classification results was achieved through the union of HydraNet segmentation and class activation maps generated using WormNet. The class activation maps analyses combined with body part segmentation in such tandem fashion allowed us to extract features responsible for lifespan prediction. Finally, using higher resolution segmented version of the *C. elegans* images, we verified our results in a higher expressive capacity residual CNN InceptionV3 accompanied by manual interpretation.

## Results

The time-lapse data for 734 *C. elegans* captured from day 1 of adulthood till death were used to develop our prototype platform (8, 14). To develop an approach for automated interpretability of these images we addressed a problem of segmenting the worms from their background, as well as, distinguishing worms’ morphological parts (Figure 1). For this, we have manually annotated 130 images of adult worms with masks for anterior, mid-body, posterior parts of the worm and summing up to a total worm mask (Figure 1F,G). This dataset was then split into the train (90) and test (40) fractions based on the dataset ID of an individual worm to ensure that individual worm features would not leak to the test hold-out. First, to address the total worm segmentation problem we have constructed a relatively shallow architecture akin to U-Net (22) accompanied with a sigmoid head for binary classification. For clarity, the encoding and decoding parts of U-Net are shown on the Figure 1A as α and β. The raw images were scaled to 96×96 pixels for computational efficiency. We used Dice loss function and monitored Jaccard index to assess the segmentation quality. On this relatively simple segmentation problem Jaccard index reached 0.97 on both train and test fractions (Figure 1A,B). Next, to extend this approach to segmentation of individual body parts of *C. elegans* we have reformulated the problem as a multi-class segmentation with one-hot encoded masks and similar U-Net-like architecture (Figure 1C,I). Unsurprisingly, since a multi-class classification is a harder problem, this led to a worse performance of 0.92 and 0.91 Jaccard index on train and test fraction respectively suggesting a mild overfit.

**Fig. 1.**
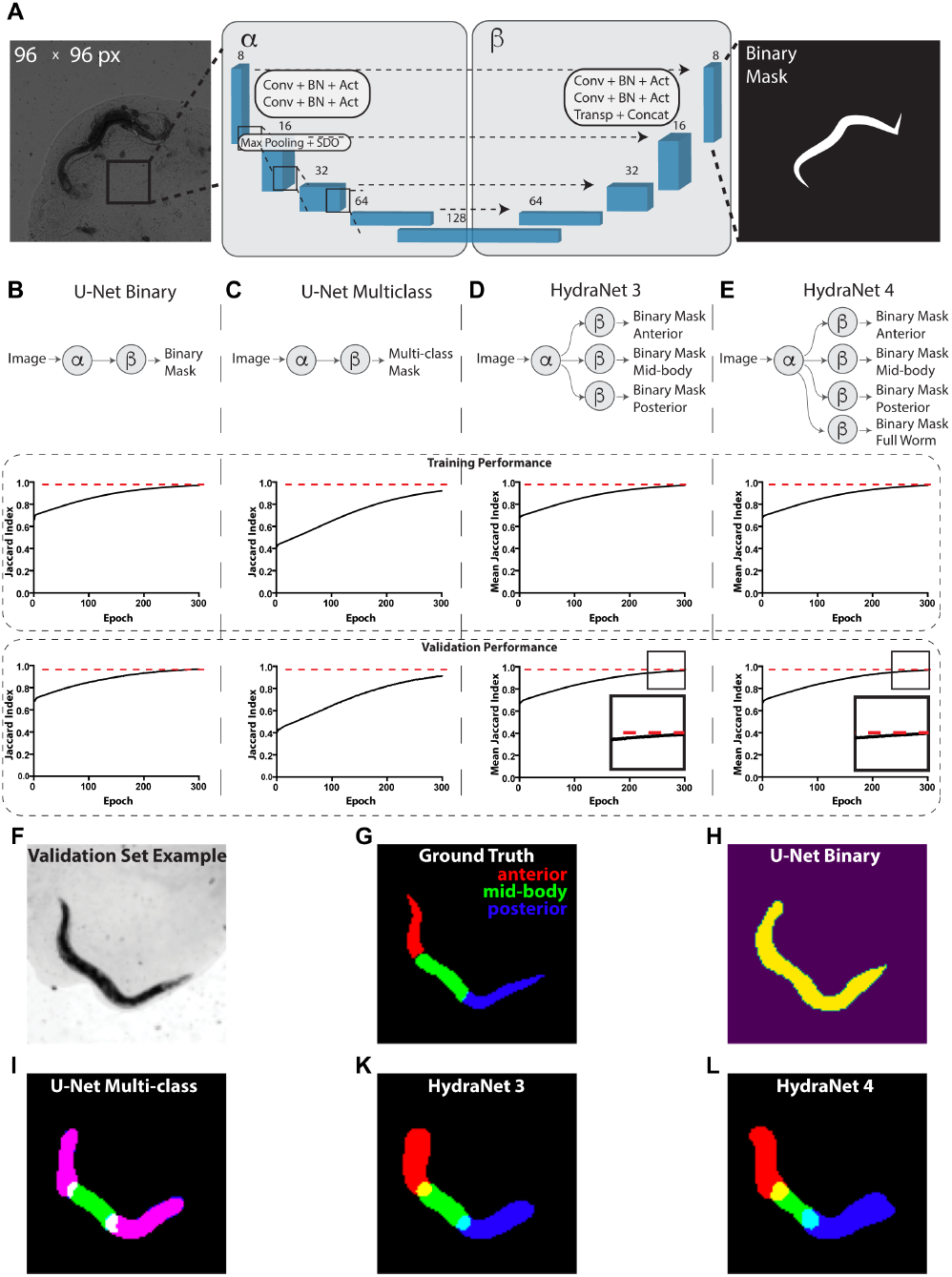
Devising worm body parts segmentation strategy. (**A**) Schematic depiction of the U-Net architecture adopted from (22). Here, transmission light micrograph of *C. elegans* used as input is depicted on a left-hand side. The reference size of the field-of-view is 580.5 μm by 580.5 μm sized to 96×96 pixels. A schematic depiction of a binary mask used as output is depicted on a right-hand side. The displayed numbers correspond to the number of filters in convolutional (Conv), batch normalisation (BN) and activation (Act) layers. Max polling layers were combined with spatial dropout (SDO). Arrows correspond to skip connections from encoder to the mirroring decoder layer where a new layer is a result of concatenation (Concat) of a layer from the encoder part to the transposed convolutional layer (Transp) from the decoder part. For illustration purposes, parts of architecture were grouped into left (*α*) and right (*β*) parts. (**B**) Schematic depiction of the binary classification U-Net architecture variant. (**C**) Schematic depiction of the multi-class classification U-Net architecture variant. (**D**) Schematic depiction of the HydraNet 3 architecture variant (**E**) Schematic depiction of the HydraNet 4 architecture variant (B-E) Here *α* is the left and *β* is the right part of the architecture in A. Graphs below show training and validation segmentation performance of the network measured as Jaccard Index for each training epoch. (**F**) Test set input data example. (**G**) Ground truth of *C. elegans* body parts segmentation example. (**H**) Output example of binary classification U-Net on the test data. (**I**) Output example of multi-class U-Net on the test data. Here, red and blue colored masks overlap making anterior and posterior parts appear magenta. (**K**) Output example of HydraNet 3 on the test data. (**L**) Output example of HydraNet 4 on the test data.

Remarkably, one aspect multi-class U-Net did not perform well was distinguishing anterior and posterior parts of the worm which led to generating overlapping masks (Figure 1I). To circumvent this limitation, we have designed an alternative architecture using U-Net α and β parts, with multiple β parts dedicated each for its own binary segmentation problem (Figure 1D,E), which we called HydraNet. Such approach creates a jointly trained architecture with common input layers and layers dedicated for each of the morphological parts of the worm, allowing to have an end-to-end model, while solving a simpler binary classification p roblem. HydraNet3 was equipped with 3 β parts dedicated to the anterior, mid-body, and posterior parts of the worm body. HydraNet4, in turn, was equipped 4 β parts dedicated to the anterior, mid-body, posterior parts as well as the whole worm body. To estimate joint performance of HydraNet we measured Jaccard index for each β parts individually and finally evaluated the average Jaccard index. Remarkably, both HydraNet3 and HydraNet4 achieved the average Jaccard index 0.97 on both the train and test fractions demonstrating good generalisation (Figure 1D,E,K,L). Noteworthy, HydraNet4 achieved conversion earlier than HydraNet3 (Figure 1D,E insets) suggesting a potential positive effect from accompanying the architecture with a more general semantic class.

Next, to obtain classifiers for *C. elegans* movement or lifespan, we split all 734 worms into 2 total movement amount classes: low or high movement estimated as motility above or below average distance crawled during the life-time; and 2 lifespan classes: ‘short-lived’ with lifespan 7 days or less, and ‘long-lived’ with lifespan 8 days and more. The task was to predict classes based on day 1, day 2 or day 3 images. As the dataset is relatively small, the use of high expressive capacity architectures could lead to overfitting. Therefore, we designed a relatively shallow CNN we called WormNet. This architecture consisted of 5 convolutional layers, each followed by a max pooling layer. Dropout and batch normalisation were implemented for each convolutional layer in the neural network to improve generalisation. The last max pooling layer was flattened and attached to a fully connected layer followed by a softmax layer. We used binary cross-entropy as a loss function. All the layers, except the latter one, used rectified linear unit (ReLU) as an activation function (Figure 2A). WormNet was used to obtain both movement and lifespan classifiers (Figures 2 and 3).

**Fig. 2.**
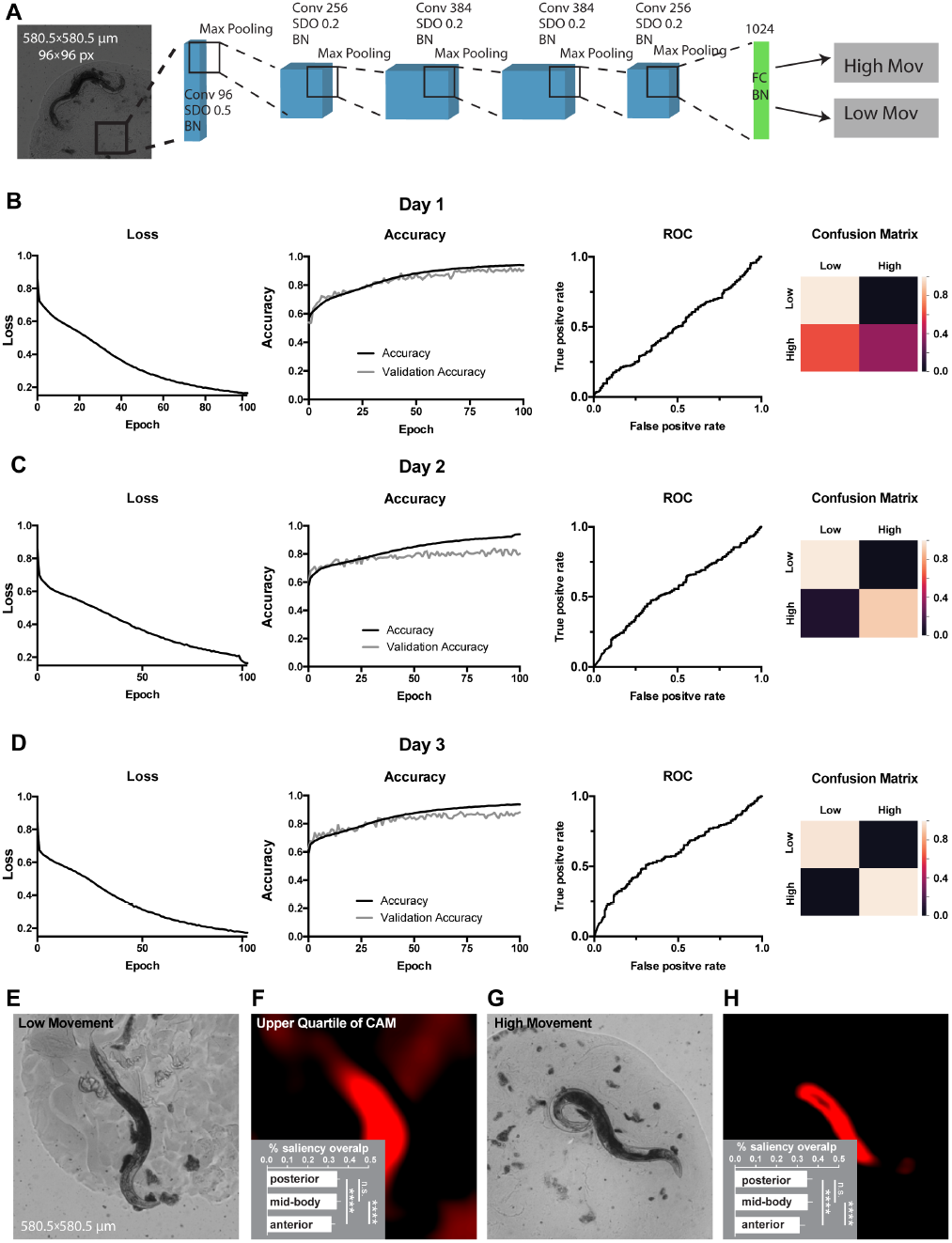
Classification of movement from end-point *C. elegans* micrographs accompanied by the by-design-interpretation based on segmentation and saliency union. (**A**) Schematic depiction of the WormNet architecture. Numbers correspond to the number of filters in convolutional (Conv), fully connected (FC), batch normalisation (BN) and activation (Act) layers. Max polling layers were combined with spatial dropout (SDO). (B-D) End-point day 1, 2 and 3 (respectively) micrographs classification loss (cost function), accuracy, receiver operating characteristic (ROC) curve, and confusion matrix. Training and test (validation) holdouts are depicted as black and light-grey lines respectively. (**E**) Low movement test micrograph example. (**F**) Upper quartile of saliency through class activation map (CAM) from image in E accompanied by the quantified by-design-interpretation using HydraNet 4 and CAM union (% saliency overlap). One-way ANOVA with Tukey’s HSD correction. Mean ± SEM, p-value < 0.0001. (**G**) High movement test micrograph example. (**H**) Upper quartile of saliency through class activation map (CAM) from image in G accompanied by the quantified by-design-interpretation using HydraNet 4 and CAM union (% saliency overlap). One-way ANOVA with Tukey’s HSD correction. Mean ± SEM, p-value < 0.0001. Here, the reference size of the field-of-view is 580.5 μm by 580.5 μm.

**Fig. 3.**
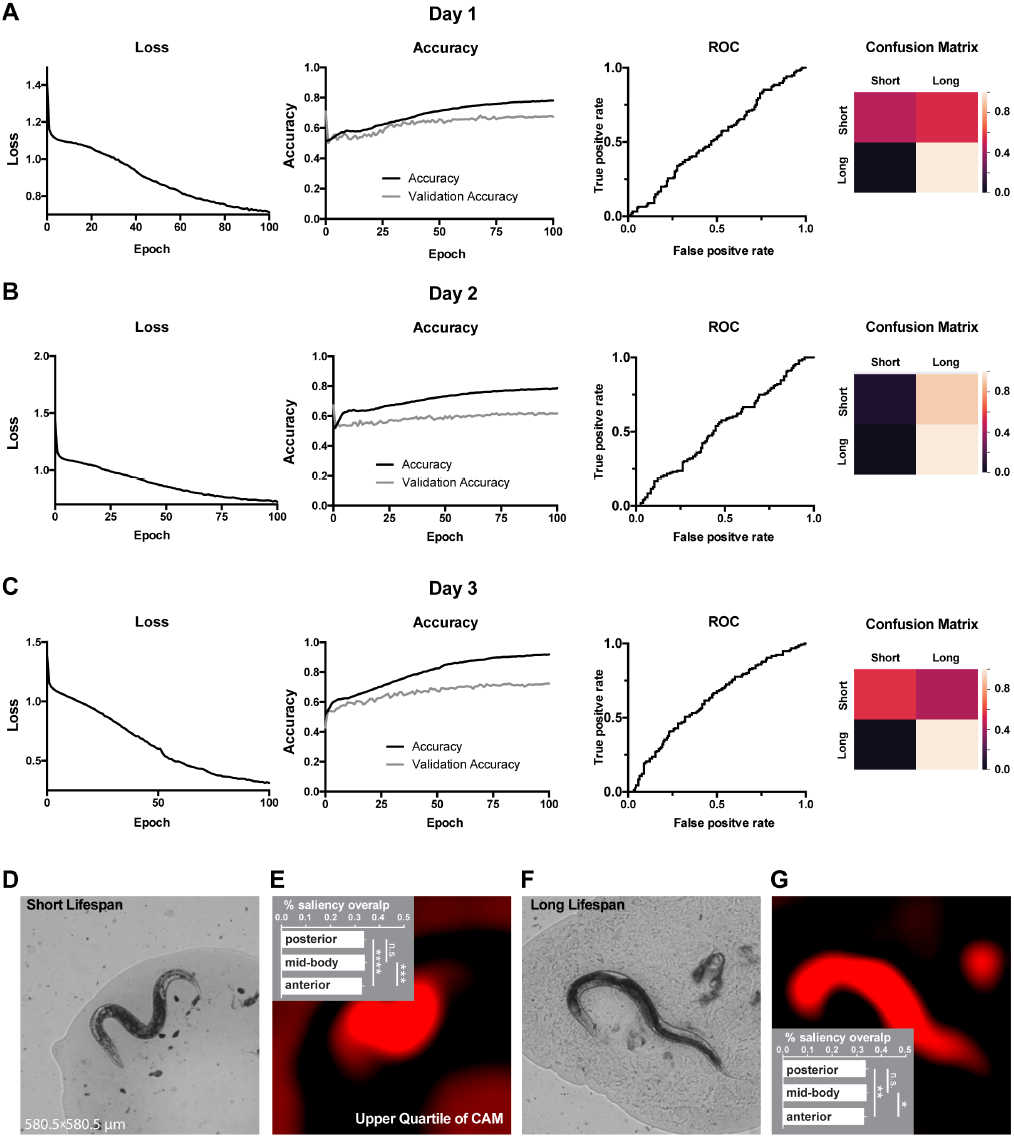
Classification of lifespan from end-point *C. elegans* micrographs accompanied by by-design-interpretation based on segmentation and saliency union. (A-C) End-point day 1, 2 and 3 (respectively) micrographs classification loss (cost function), accuracy, receiver operating characteristic (ROC) curve, and confusion matrix. Training and test (validation) holdouts are depicted as black and light-grey lines respectively. (**D**) Short lifespan test micrograph example. (**E**) Upper quartile of saliency through class activation map (CAM) from image in D accompanied by the quantified by-design-interpretation using HydraNet 4 and CAM union (% saliency overlap). One-way ANOVA with Tukey’s HSD correction. Mean ± SEM, ***p-value < 0.001, ****p-value < 0.0001. (**F**) Long lifespan test micrograph example. (**G**) Upper quartile of saliency through class activation map (CAM) from image in F accompanied by the quantified by-design-interpretation using HydraNet 4 and CAM union (% saliency overlap). One-way ANOVA with Tukey’s HSD correction. Mean ± SEM, *p-value < 0.05, **p-value < 0.01. . Here, the reference size of the field-of-view is 580.5 μm by 580.5 μm.

The WormNet showed good performance on total movement classification reaching 88% accuracy (precision 0.86, recall 0.86, area under curve for receiver operating characteristic - AUC ROC - was 0.56) on the test dataset for the day 3 adults fraction. The performance for the day 1 and day 2 images were slightly lower (Figure 2B-D) with ROC AUC of 0.51 and 0.55 respectively. To assess which body part might be responsible for the WormNet decision making, using our tandem segmentation-classification approach we have obtained CAMs for a low movement class worm (Figure 2E,F) and a high movement worm (Figure 2G,H) from WormNet. Next, each image was segmented using HydraNet4 and the union of WormNet upper quartile CAM with morphological part segmentation from HydraNet4 was obtained. For interpretation purposes we have computed the percentage of CAMs belonging to a respective morphological segment for each respective worm belonging to high or low movement class. Furthermore, we assessed significance of this by-design interpretation using one-way ANOVA with Tukey’s honest significant difference (HSD) correction (Figure 2F – low movement worms, Figure 2H – high movement worms). The comparison suggested that the anterior part was covered significantly less (31%) than mid-body (34%) and posterior parts (34%) for both low and high movement worms. There was no significant difference between the mid-body and the posterior part of the body.

Next, we used WormNet to classify long and short-lived worms. Similarly to movement classification, the WormNet performed better on day 3 adults sample reaching accuracy of 72% (precision 0.73, recall 0.71, AUC ROC 0.61) on the test dataset, as compared to AUC ROC of 0.53 and 0.52 for day 2 and 1 respectively. The confusion matrix analysis suggested that the CNN underperformed in short-lived worms classifying (Fig. 3A). Next, we have interpreted the classifier using the tandem of HydraNet4 and WormNet accompanied by one-way ANOVA statistical test. In the case of lifespan classification, by-design interpretation suggested that at 32% the anterior part was significantly less pronounced in CAMs comparing to the mid-body and the posterior part (Figure 3E - short lifespan, 3G – long lifespan). This difference was less significant for long lifespan than for short lifespan. There was no significant difference between the mid-body and the posterior part.

To verify these findings in an independent manner we have trained another lifespan classifier u sing t he r esidual InceptionV3 architecture (31) accompanied by a manual interpretation (Fig. 4). Furthermore, in this case to ensure high resolution of the CAMs instead of scaling to 96×96 pixels, the full resolution 900×900 images cropped to 800×800 pixels (516×516 μm) were used. As a much higher expressive capacity CNN, InceptionV3 was prone to overfitting on our relatively small dataset (Figure 4C,D). To circumvent this, we have implemented early stopping during training. Additionally, we segmented the worms from their background ensuring InceptionV3 is presented only with the relevant part of the image. InceptionV3 performed similarly to WormNet with the accuracy reaching 70% on the test dataset for lifespan classification (Figure 4A). Consistently with the tandem HydraNet4-WormNet approach to interpretation, in the case of the manual interpretation, the anterior part of the worm was highlighted by the InceptionV3 CAM less frequently. Importantly, however, due to the higher resolution of the input images, the CAMs now localised the body parts much better, allowing to assign a body part as a possible discriminator in each case (Figure 4B). Interestingly, the distribution of the body parts highlighted by CAM’s analysis demonstrates that posterior part is more important for long-living worms’ classification, suggesting that the features predicting longevity could be located in the posterior part of the worm body.

**Fig. 4.**
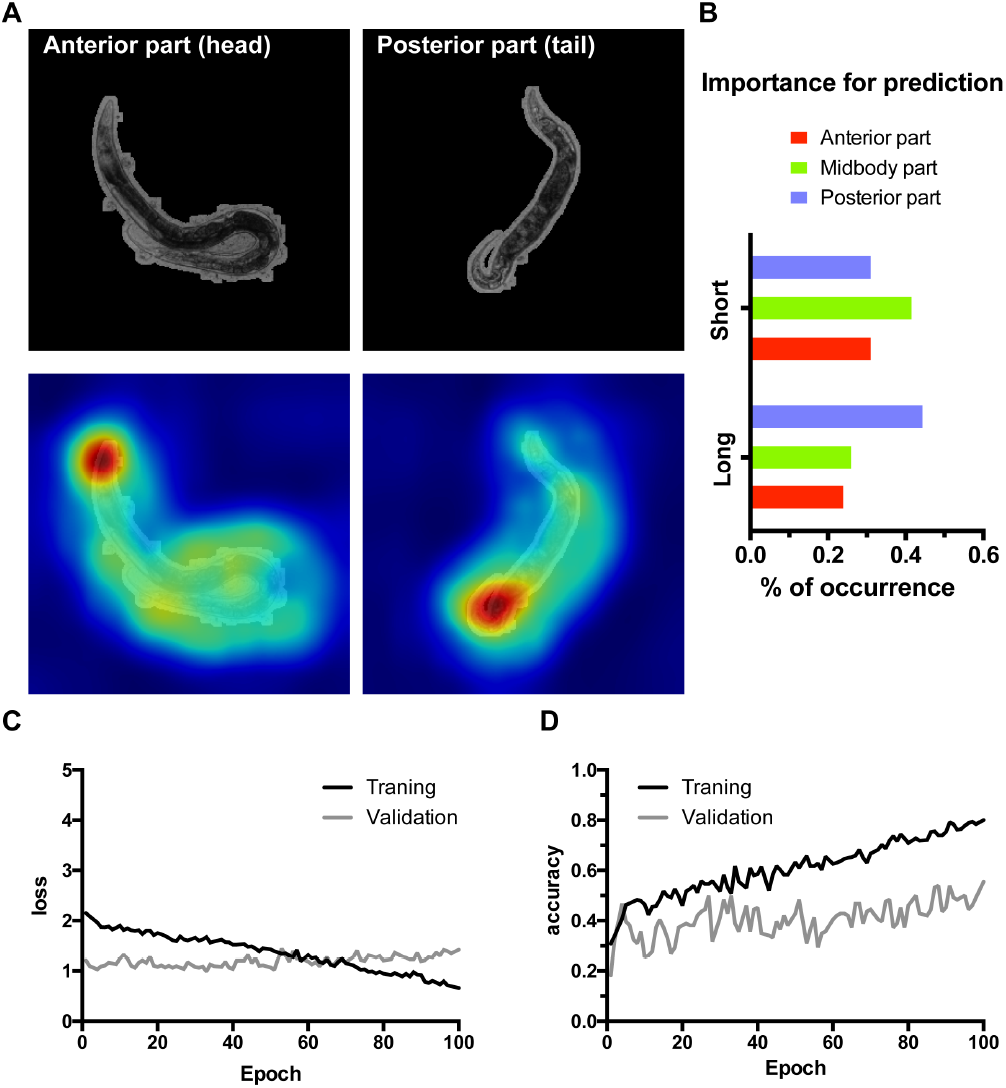
Class activation maps (CAMs) of the InceptionV3 network overfitted on high resolution images allow localising lifespan-related regions. (**A**) Examples of the class activation maps pointing to anterior (left) or posterior (right) parts of a worm. Here 516 by 516 μm area was represented by the 800×800 px input image for higher resolution input. (**B**) *C. elegans* body parts importance for class prediction measured as the percent of occurrence of body parts highlighted on the CAMs. (**C - D**) Loss and accuracy training statistics for InceptionV3 network.

## Discussion

Despite *C. elegans* is a classical model in ageing research with more than 4000 papers published up to date, and the progress in robotics, the process of measuring *C. elegans* lifespan is still manual and laborious. However, new approaches are emerging like lifespan machine utilizing flatbed scanners to simultaneously assess the viability of large population of worms on plates (32). Another approach is Worm corals – an automated vermiculture methods allowing to track worms throughout their lifespan with much better detailed measurements (8). The detailed physiological data produced on Worm corals showed that movement, autofluorescence and textural degradation are the best predictors of lifespan. However, it remains unclear what exact morphological features reflect pathologies and determine the lifespan length. It was also found that physiological measurements before day 3 or 4 of adulthood as well as single GFP labelled biomarkers are not able to distinguish short and long-lived worms (8, 14). Nucleolar-based predictions made on day 1 adults are performed using 100x magnification on fixed worms, which is not achievable for any automated screening platform.

Here we worked with the dataset generated in Pincus lab (8, 14), and showed that the application of newly designed WormNet was able to successfully discriminate between short and long-lived worms even for images taken at day 1 or day 2; importantly, for day 3 the CNN demonstrated the best performance (Fig. 2A-C). WormNet was even better at classifying worms with high and low total movement, achieving 88% accuracy for day 1 adults (Fig. 3). We expect that generating more data and developing the CNN architecture could further improve the performance of WormNet by decreasing overfitting and bias. CNN were used before for regressing and classifying *C. elegans* lifespan (29). The authors manually classified curvy and straight postures of nematodes which allowed them to improve their accuracy, which still remained relatively low due to small sample size (913 images). Pincus dataset was also used to assess CNN ability to predict lifespan (30). As mentioned earlier, the authors segmented the worms and created straightened worm representations, which were used for CNN training (30). Increased number of samples improved the regression-based prediction of worm age. Interestingly, the authors found that worm silhouette alone has limited information for age estimation, whereas the information from background can significantly improve the accuracy, though the predictive value of background is an artifact of experimental conditions. Therefore, it might be possible the predictive accuracy of WormNet in our simulations can be partly explained by the background information. Importantly, pretraining on the body-coordinate representations improved accuracy on raw images which suggests that worm organs and texture are useful for age prediction.

In addition to lifespan or movement classification based on young adults’ images, we also aimed to find features important for the prediction. As a prototype task we decided to determine which body part – anterior, mid-body or posterior part contains features influencing lifespan length the most.

We designed HydraNet 3 and 4, new architectures based on U-Net and showed that they can successfully segment worm body parts achieving perfect Jaccard index values. Importantly, to develop a by-design interpretation approach we employed a tandem of biologically meaningful classification (lifespan and movement) yielding saliency through class activation maps (28, 33) and morphological segmentation (anterior, mid-body and posterior regions) to find which body part is useful for the classifications. Furthermore, although less resolved, findings obtained from the tandem approach were consistent with an independently trained classifier. This binary classifier was based on the InveptionV3 CNN. It was trained on 800×800 pixel full optical resolution images with worms segmented from their background and achieved results comparable to WormNet, though the model is less generalizable due to more overfitting (Figure 4). However, in the case of InceptionV3, distinct body parts could be localized on the CAMs, and the analyses suggests that features located in the posterior part of the worm might be more important for classifying long-lived worms.

This approach provides avenue to the discovery of new important age biomarkers in *C. elegans* in automated setting, given a significant increase in image resolution and usage of body-coordinate representation. Non-labelled organs like pharynx or GFP-labelled entities could be segmented using HydraNets and assessed for their lifespan predictive ability using CAM approach and WormNet. It is tempting to speculate that akin to generative adversarial networks (34), future implementations of the by-design interpretability through a tandem of segmentation and classification may be trained end-to-end and employed for routine scientific discovery. The proof-of-principle automated analytical platform will be useful for non-invasive aging biomarkers discovery, particularly in young day 1-3 adult *C. elegans*. This has a great potential to accelerate the pharmaceutical screening for anti-aging drugs. The development of the methodology will also be helpful to find and characterize new pathologies in *C. elegans* important for basic ageing research.

## ACKNOWLEDGEMENTS

We would like to thank Pincus lab for providing us with the access to the *C. elegans* dataset.

## Methods

### Code implementation

All the source code for this work was implemented in Python version 3.6, Tensorflow versions 1.9.0 or 2.3.0 (Abadi et al., 2016) and Keras version 2.2.2 (Chollet, 2018). Tensors pre-processing and manipulation was implemented in Numpy (Harris et al., 2020). Python environment was maintained using anaconda distribution.

### Model training

Training of WormNet, and fine-tuning of InceptionV3 were performed on a desktop PC equipped with Intel Core i7-8700K CPU at 3.7 GHz and 32 GB of RAM as well as GeForce 1080 Ti GPU. Training of U-Net and HydraNet were performed using Google Collaboratory cloud GPU (e.g. NVIDIA Tesla V100). Inference and analysis were performed using Google Collaboratory.

### Statistical analysis

Statistical significance was evaluated using one-way ANOVA with Tukey’s HSD correction employing GraphPad Prism software.

### Architecture design and Hyperparameters tuning

To ensure optimal performance of U-Net, WormNet, HydraNet and InceptionV3 architectures hyperparameters including, but not limited to the rate of learning, regularization through dropout coefficient were heuristically optimised. For the baseline comparison, all hyperparameters across similar architectures were kept comparable. In case of novel architectures, during the design process the expressive capacity or depth of the architecture was started at lowest applicable size and gradually increased until the point of performance convergence.

## References

1. Partridge, L., Fuentealba, M., and Kennedy, B. K. The quest to slow ageing through drug discovery. Nature Reviews Drug Discovery, 19(8):513–532, 2020.

2. Caldwell, K. A., Willicott, C. W., and Caldwell, G. A. Modeling neurodegeneration in caenorhabditis elegans. Disease Models and Mechanisms, 13(10), 2020.

3. Galimov, E. R., Pryor, R. E., Poole, S. E., Benedetto, A., Pincus, Z., and Gems, D. Coupling of rigor mortis and intestinal necrosis during c. elegans organismal death. Cell reports, 22 (10):2730–2741, 2018.

4. Ezcurra, M., Benedetto, A., Sornda, T., Gilliat, A. F., Au, C., Zhang, Q., van Schelt, S., Petrache, A. L., Wang, H., de la Guardia, Y., Bar-Nun, S., Tyler, E., Wakelam, M. J., and Gems, D. C. elegans eats its own intestine to make yolk leading to multiple senescent pathologies. Current Biology, 28(16):2544–2556, 2018.

5. Wang, H., Zhao, Y., Ezcurra, M., Benedetto, A., Gilliat, A. F., Hellberg, J., Ren, Z., Galimov, E. R., Athigapanich, T., and Girstmair, J. A parthenogenetic quasi-program causes teratoma-like tumors during aging in wild-type c. elegans. npj Aging and Mechanisms of Disease, 4(1):1–12, 2018.

6. Zhao, Y., Gilliat, A. F., Ziehm, M., Turmaine, M., Wang, H., Ezcurra, M., Yang, C., Phillips, G., McBay, D., and Zhang, W. B. Two forms of death in ageing caenorhabditis elegans. Nature communications, 8(1):1–8, 2017.

7. LeCun, Y. and Bengio, Y. Convolutional networks for images, speech, and time series. The handbook of brain theory and neural networks, 3361(10):1995, 1995.

8. Zhang, W. B., Sinha, D. B., Pittman, W. E., Hvatum, E., Stroustrup, N., and Pincus, Z. Extended twilight among isogenic c. elegans causes a disproportionate scaling between lifespan and health. Cell Syst, 3(4):333–345 e4, 2016.

9. Pickett, C., Dietrich, N., Chen, J., Xiong, C., and Kornfeld, K. Mated progeny production is a biomarker of aging in caenorhabditis elegans. G3: Genes, Genomes, Genetics, 3(12): 2219–2232, 2013.

10. Hahm, J.-H., Kim, S., DiLoreto, R., Shi, C., Lee, S.-J. V., Murphy, C. T., and Nam, H. G. C. elegans maximum velocity correlates with healthspan and is maintained in worms with an insulin receptor mutation. Nature communications, 6(1):1–7, 2015.

11. Hsu, A.-L., Feng, Z., Hsieh, M.-Y., and Xu, X. S. Identification by machine vision of the rate of motor activity decline as a lifespan predictor in c. elegans. Neurobiology of aging, 30(9): 1498–1503, 2009.

12. Rea, S. L., Wu, D., Cypser, J. R., Vaupel, J. W., and Johnson, T. E. A stress-sensitive reporter predicts longevity in isogenic populations of caenorhabditis elegans. Nature genetics, 37(8):894–898, 2005.

13. Sánchez-Blanco, A. and Kim, S. K. Variable pathogenicity determines individual lifespan in caenorhabditis elegans. PLoS genetics, 7(4):e1002047, 2011.

14. Pincus, Z., Smith-Vikos, T., and Slack, F. J. Microrna predictors of longevity in caenorhabditis elegans. PLoS Genet, 7(9):e1002306, 2011.

15. Tiku, V., Jain, C., Raz, Y., Nakamura, S., Heestand, B., Liu, W., Späth, M., Suchiman, H. E. D., Müller, R.-U., and Slagboom, P. E. Small nucleoli are a cellular hallmark of longevity. Nature communications, 8(1):1–9, 2017.

16. Eckley, D. M., Rahimi, S., Mantilla, S., Orlov, N. V., Coletta, C. E., Wilson, M. A., Iser, W. B., Delaney, J. D., Zhang, Y., and Wood, W. Molecular characterization of the transition to mid-life in caenorhabditis elegans. Age, 35(3):689–703, 2013. ISSN 0161-9152.

17. Cireşan, D., Meier, U., and Schmidhuber, J. Multi-column deep neural networks for image classification. arXiv preprint arXiv:1202.2745, 2012.

18. Krizhevsky, A., Sutskever, I., and Hinton, G. E. Imagenet classification with deep convolutional neural networks. Advances in neural information processing systems, 25:1097–1105, 2012.

19. Russakovsky, O., Deng, J., Su, H., Krause, J., Satheesh, S., Ma, S., Huang, Z., Karpathy, A., Khosla, A., and Bernstein, M. Imagenet large scale visual recognition challenge. International Journal of Computer Vision, 115(3):211–252, 2015.

20. Hanif, M. S. and Bilal, M. Competitive residual neural network for image classification. ICT Express, 6(1):28–37, 2020.

21. He, K., Zhang, X., Ren, S., and Sun, J. Deep residual learning for image recognition. In Proceedings of the IEEE conference on computer vision and pattern recognition, pages 770–778.

22. Ronneberger, O., Fischer, P., and Brox, T. U-net: Convolutional networks for biomedical image segmentation. In International Conference on Medical image computing and computer-assisted intervention, pages 234–241. Springer.

23. Du, G., Cao, X., Liang, J., Chen, X., and Zhan, Y. Medical image segmentation based on u-net: A review. Journal of Imaging Science and Technology, 64(2):20508–1–20508–12, 2020.

24. Mamoshina, P., Vieira, A., Putin, E., and Zhavoronkov, A. Applications of deep learning in biomedicine. Molecular pharmaceutics, 13(5):1445–1454, 2016.

25. Brahimi, M., Arsenovic, M., Laraba, S., Sladojevic, S., Boukhalfa, K., and Moussaoui, A. Deep learning for plant diseases: detection and saliency map visualisation, pages 93–117. Springer, 2018.

26. Fisch, D., Yakimovich, A., Clough, B., Wright, J., Bunyan, M., Howell, M., Mercer, J., and Frickel, E. Defining host–pathogen interactions employing an artificial intelligence workflow. Elife, 8:e40560, 2019.

27. Zeiler, M. D. and Fergus, R. Visualizing and understanding convolutional networks. In European conference on computer vision, pages 818–833. Springer.

28. Zhou, B., Khosla, A., Lapedriza, A., Oliva, A., and Torralba, A. Learning deep features for discriminative localization. In Proceedings of the IEEE Conference on Computer Vision and Pattern Recognition, pages 2921–2929.

29. Lin, J.-L., Kuo, W.-L., Huang, Y.-H., Jong, T.-L., Hsu, A.-L., and Hsu, W.-H. Using convolutional neural networks to measure the physiological age of caenorhabditis elegans. IEEE/ACM transactions on computational biology and bioinformatics, 2020.

30. Wang, L., Kong, S., Pincus, Z., and Fowlkes, C. Celeganser: Automated analysis of nematode morphology and age. In Proceedings of the IEEE/CVF Conference on Computer Vision and Pattern Recognition Workshops, pages 968–969.

31. Szegedy, C., Vanhoucke, V., Ioffe, S., Shlens, J., and Wojna, Z. Rethinking the inception architecture for computer vision. In Proceedings of the IEEE conference on computer vision and pattern recognition, pages 2818–2826.

32. Stroustrup, N., Ulmschneider, B. E., Nash, Z. M., López-Moyado, I. F., Apfeld, J., and Fontana, W. The caenorhabditis elegans lifespan machine. Nature methods, 10(7):665–670, 2013.

33. Sun, K. H., Huh, H., Tama, B. A., Lee, S. Y., Jung, J. H., and Lee, S. Vision-based fault diagnostics using explainable deep learning with class activation maps. IEEE Access, 8: 129169–129179, 2020.

34. Goodfellow, I., Pouget-Abadie, J., Mirza, M., Xu, B., Warde-Farley, D., Ozair, S., Courville, A., and Bengio, Y. Generative adversarial nets. In Advances in neural information processing systems, pages 2672–2680.

